# Inclusion of a GaAs detector model in the Photon Counting Toolkit software for the study of breast imaging systems

**DOI:** 10.1101/2022.06.10.495610

**Authors:** Bahaa Ghammraoui, Katsuyuki Taguchi, Stephen J. Glick

## Abstract

We present an upgraded version of the Photon Counting Toolkit ([PcTK](https://pctk.jhu.edu/)), a publicly available MATLAB tool for the simulation of semiconductor-based photon counting detectors (PCD), which has been extended and validated to account for gallium arsenide (GaAs)-based PCD(s). The modified PcTK version was validated by performing simulations and acquiring experimental data for three different cases. The LAMBDA 60 K module planar detector (X-Spectrum GmbH, Germany) based on the Medipix3 ASIC technology was used in all cases. This detector has a 500-*µ*m thick GaAs sensor and a 256 × 256-pixel array 55 *µ*m pixel size. The first validation was a comparison between simulated and measured spectra from a ^109^Cd radionuclide source. In the second validation study, experiments and simulations of mammography spectra were conducted to observe the performance of the GaAs version of the PcTK with polychromatic radiation used in conventional x-ray imaging systems. The third validation study used single event analysis to validate the spatio-energetic model of the extended PcTK version. Overall, the software provided a good agreement between simulated and experimental data, validating the accuracy of the GaAs model. The software could be an attractive tool for accurate simulation of breast imaging modalities relying on photon counting detectors and therefore could assist in their characterization and optimization.

## Introduction

Clinical x-ray imaging equipment has been operating in energy integrating mode for more than a century. During the last few years, we have been on the verge of a paradigm shift in clinical x-ray detector technology, with future detectors enabling photon counting and energy discrimination. These Photon Counting Detector (PCD) systems have several advantages, particularly in breast imaging [1]. Previous studies have shown that photon counting imaging systems can reduce electronic noise and beam-hardening, improve contrast-to-noise ratio through energy weighting, improve radiation dose efficiency, and could enable extremely high spatial resolution. Furthermore, simultaneous multi-energy data acquisition would allow differentiation among multiple contrast agents [2]. It is well known that most of the major computed tomography (CT) companies have been developing prototype PCD-CT systems recently, and last year the FDA cleared the world’s first PCD-CT scanner, the Siemens Naeotom Alpha. This current rapid development of PCD spectroscopic imaging systems is made possible by recent technological developments in integrated circuits for reading signals from solid state detectors, and advances in the capabilities of the maximum count rate, as well as the lower cost and improved reliability of larger semiconductor boards.

However, several technical challenges still must be overcome in order to realize the full potential of PCD. These include charge sharing, pulse pile-up, characteristic escape peaks, Compton scattering, and weighting potential cross talk, which can severely distort measured spectra and spatial resolution in such pixelated detectors [3, 4]. It is widely recognized that computer simulation codes and models can help, and they represent key components of the development, optimization and evaluation of such imaging systems. To date, only one publicly available code has been developed and dedicated to modeling semiconductor-based PCDs. This code is part of the Photon Counting Toolkit (PcTK) software [5], which was developed mainly to model cadmium telluride (CdTe)-based PCDs. Other semiconductor-based x-ray PCD candidates, including silicon (Si) and gallium arsenide (GaAs), are under investigation and development and are not yet modeled in the PcTK. GaAs, specifically, is promising for breast imaging, since high quantum efficiencies are easily achievable within the mammography energy range (12 to 45 KeV). Furthermore, its characteristic K-edges lie below the mammography-relevant energies (9, 10, and 12 keV), minimizing the likelihood of fluorescence x-rays escaping the pixels.

In a recent study, Ghammraoui et al. showed that GaAs spectral mammography provides slightly improved or equivalent performance versus commercial mammography systems equipped with energy integrated detectors [6]. In that study, commercial mammography system exposure irradiation conditions were used, and no attempt was made to optimize parameters for the GaAs detector. Such an optimization study could easily be conducted using a reliable experimentally validated GaAs PCD simulation code to complement experimental work. Appropriate optimization efforts could reduce development time and costs.

In this work, we present an extension to the PcTK that enables realistic GaAs PCD simulation. We first describe the upgraded simulation parameters needed to model GaAs and then provide information on the three experimental procedures used for parameter optimization and code validation. Such publicly available code with GaAs simulation could promote PCD research in mammography, CT and digital breast tomosynthesis (DBT) modalities.

## Materials and Methods

### Detector model

The PcTK code was extended to account for simulation of a GaAs semiconductor with an atomic ratio As/Ga of 1. The extended version uses the same design concept that was used in PcTK version 3.2, described in a previous study [5]. Each element is identical to version 3.2 except for the inclusion of the GaAs interaction probabilities, with several assumptions described below, as it is meant for the energy range pertinent to breast imaging.

The GaAs model accounts for the probabilities of the different interaction phenomena in a GaAs sensor in different ways. We ignored Compton scattering interactions, since there is a low probability of these in the mammography energy range. Therefore, given an x-ray interaction absorbed within the detector, the conditional probabilities at energy E of photoelectric effect are assumed to be equal to 1. The conditional probability of K-shell void given photoelectric effect for both Ga and As was assumed to be 88% [7]. We also used experimentally measured conditional probability of fluorescence x-ray emission, given the K-shell void present (*W*_*k*_=0.528 for Ga and *W*_*k*_=0.589 for As) [8], and assumed fixed yields of 0.9 and 0.1 for K-alpha and K-beta x-ray fluorescence [9]. Finally, the mean travel distances of Ga and As fluorescence x-rays were assumed to be 42*µ*m and 16*µ*m, respectively [10]. Other parameters, such as detector thickness, pixel size *d*_0_, charge cloud radius *r*_0_ and the electronic noise parameter *σ*_*e*_ can vary according to detector specifications, and thus are to be input by the user.

### Validation

The modified PcTK version was validated by performing simulations and experimental data for three different cases. The LAMBDA 60 K module planar detector (X-Spectrum GmbH, Germany) based on the Medipix3 ASIC technology was modeled for simulations and used for validations using experimental measurements. This detector has a 500-*µ*m-thick GaAs sensor and a 256 *×* 256-pixel array 55 *µ*m pixel size.

### Optimization study

The first experiment aimed for both validation and to obtain the optimum *r*_0_ and *σ*_*e*_ model parameters for simulating the LAMBDA 60 K module planar detector used in these studies. Experimentally, the spectrum emitted from a ^109^Cd sealed source was measured by sweeping the energy threshold from 5 to 30 keV with a decrement of 2 keV, and the sweeping process was repeated several times to acquire the mean spectra with lower variability. Detector pixel spectra were calculated by differentiation between successive images. The spectrum also was averaged over 400 randomly selected pixels. In addition, simulations were performed using different *r*_0_ and *σ*_*e*_ model parameters assuming monochromatic radiation of 22 keV (primary emission from ^109^Cd). The simulated *r*_0_ values ranged from 5 to 15 *µ*m with 1 *µ*m spacing; and *σ*_*e*_ values ranged from 0.5 to 2.5 keV with 0.1 keV spacing. The simulated detector pixel size and thickness were set to 55 *µ*m and 500-*µ*m, respectively. As a figure-of-merit, the normalized cross correlation factors between the measured and simulated spectra were used. The optimal parameter values that maximize the correlation factor were selected for the further measurements and were found to be 11 *µ*m and 2.1 keV for *r*_0_ and *σ*_*e*_, respectively as shown in Figure 1 (a). It can be seen in Figure 1(b) that the experimental spectrum was correctly reproduced in the simulation by the updated tool using these optimum parameter values for *r*_0_ and *σ*_*e*_.

**Figure 1.**
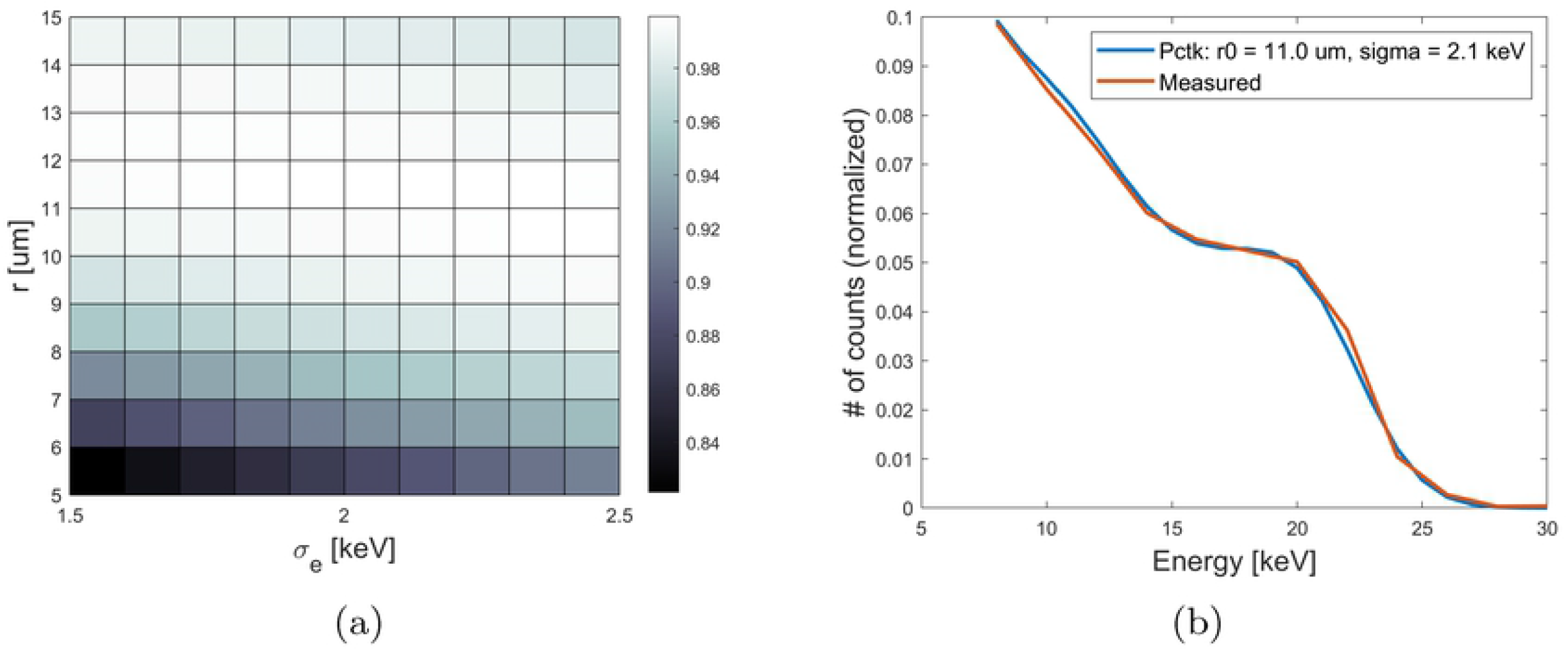
(a) shows the normalized cross correlation factors between the measured ^109^Cd spectrum and 100 simulated spectra with different *r*_0_ and *σ*_*e*_ model parameters. (b) comparison between the experimental and simulated spectra when using 11 *µ*m and 2.1 keV for *r*_0_ and *σ*_*e*_, respectively.

### Experimental validation using a polychromatic x-ray spectra

For validation in the second experiment, experimental and simulated mammography spectra were used to observe the performance of the GaAs version of the PcTK with polychromatic radiation used in conventional x-ray imaging systems. Polychromatic spectra emitted from an industrial X-ray tube with a tungsten target which operated at 30, 35 and 40 kVp, and 2 mA were measured and compared to simulated spectra using the GaAs version of the PcTK. The detector was placed 1 m away from the source, and 2.7 mm of aluminum filtration was placed in front of the source to mimic the mammography x-ray spectra transmitted through a breast. The spectra were measured by sweeping the energy threshold from 5 to 49 keV with a decrement of 2 keV, and the sweeping process was repeated several times to acquire the average and obtain better statistics. Final spectra were calculated by differentiation between successive images. The same spectra were also measured using the XR100T-CdTe Amptek (Amptek, Bedford, MA, USA) detector and corrected for spectral distortion using the simulated system response matrix described by Ghammraoui et al. [11]. The corrected spectra from the Amptek detector were the inputs to the GaAs version of the PcTK. As mentioned earlier, the optimal parameter values of *r*_0_ and *σ*_*e*_ found in the first study were used in the simulation. Figure 2 shows the simulated spectra using the PcTK and the measured spectra using the GaAs LAMBDA detector. The plots confirm good performance of the algorithm with polychromatic radiation. However, a slightly worse performance was observed for 40 and 45 kVp spectra, especially at the higher energy range. The origin of this small disagreement can be explained by different factors, including inaccurate inputs in the incident spectra, which were estimated from the Amptek one pixel detector measurements followed by the spectral analysis correction, homogeneity in the pixels spectral response, and inaccurate inputs for *r*_0_ and *σ*_*e*_.

**Figure 2.**
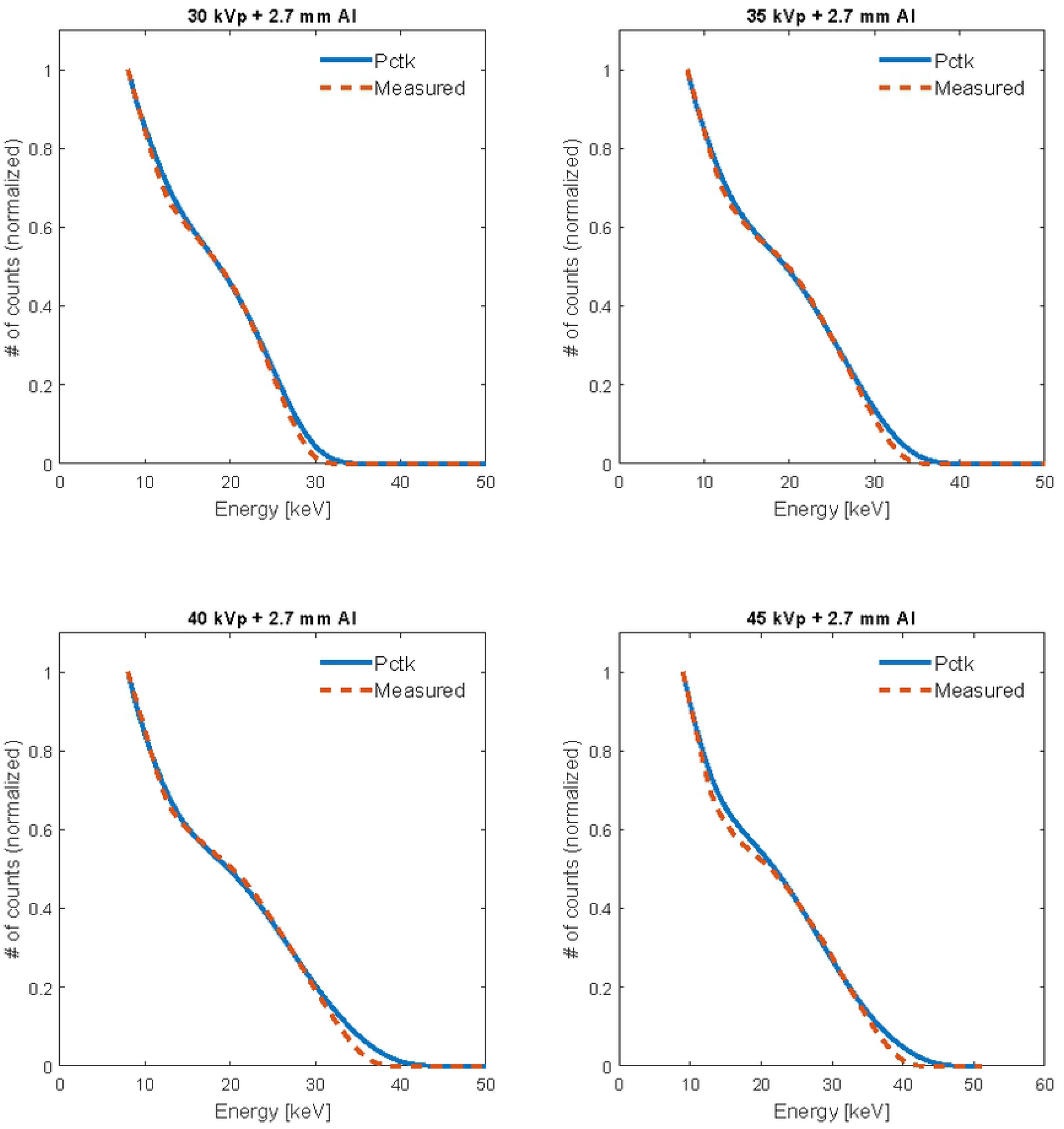
Comparisons between simulated and measured spectra with incident tungsten target X-ray spectra.

### Experimental validation of the spatio-energetic model using single events analysis

The third experiment involved validating the spatio-energetic model of the GaAs version of the PcTK using single events analysis [12]. Single events analysis was performed by exposing the detector to low fluence in order to generate point-like images where only clusters from one absorbed x-ray were present, as shown in Figure **??**. This was done by placing the detector 7 m away from the source and reducing the x-ray generator-operated current and voltage to 0.1 mA and 35 kVp, respectively. For each energy threshold acquisition, 1,800 frames were recorded, and the frame time was set to 1 ms. The analysis was performed on each frame separately to account for the number of events in which: (1) *N*_*single*_(*E*_*threshold*_) we had only one separate pixel activated and (2) *N*_*multiple*_ we had multiple adjacent pixels activated. The single events experiment was simulated using a 3×3 pixels detector and only irradiating the central pixel. Such analysis can be used to estimate the spectra recorded by the central pixel (irradiated), *I*_*central*_(*E*_*threshold*_) and its neighboring pixels *I*_*neighbor*_(*E*_*threshold*_) separately.

Experimentally, *I*_*central*_(*E*) and *I*_*neighbor*_(*E*) can also be estimated from the measured *N*_*single*_(*E*) and *N*_*multiple*_(*E*) values, using the following equations:

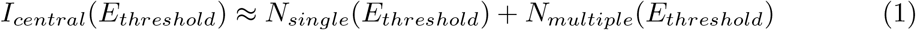

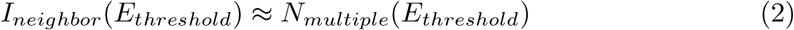

Figure 3 shows a comparison between the estimated and simulated *I*_*central*_(*E*) and *I*_*neighbor*_(*E*) spectra (*≈* differentiation between two successive *I*_*central*_(*E*_*threshold*_) and *I*_*neighbor*_(*E*_*threshold*_), respectively). As can be observed, the measured spectra are correctly reproduced using the GaAs version of the PcTK. Although some disagreements could be observed, especially at the higher ends of the spectra, the plots confirm good overall performance of the model used, and these disagreements can be considered negligible. Better performance could be possibly achieved by tuning the *r*_0_ and *σ*_*e*_ model parameters.

**Figure 3.**
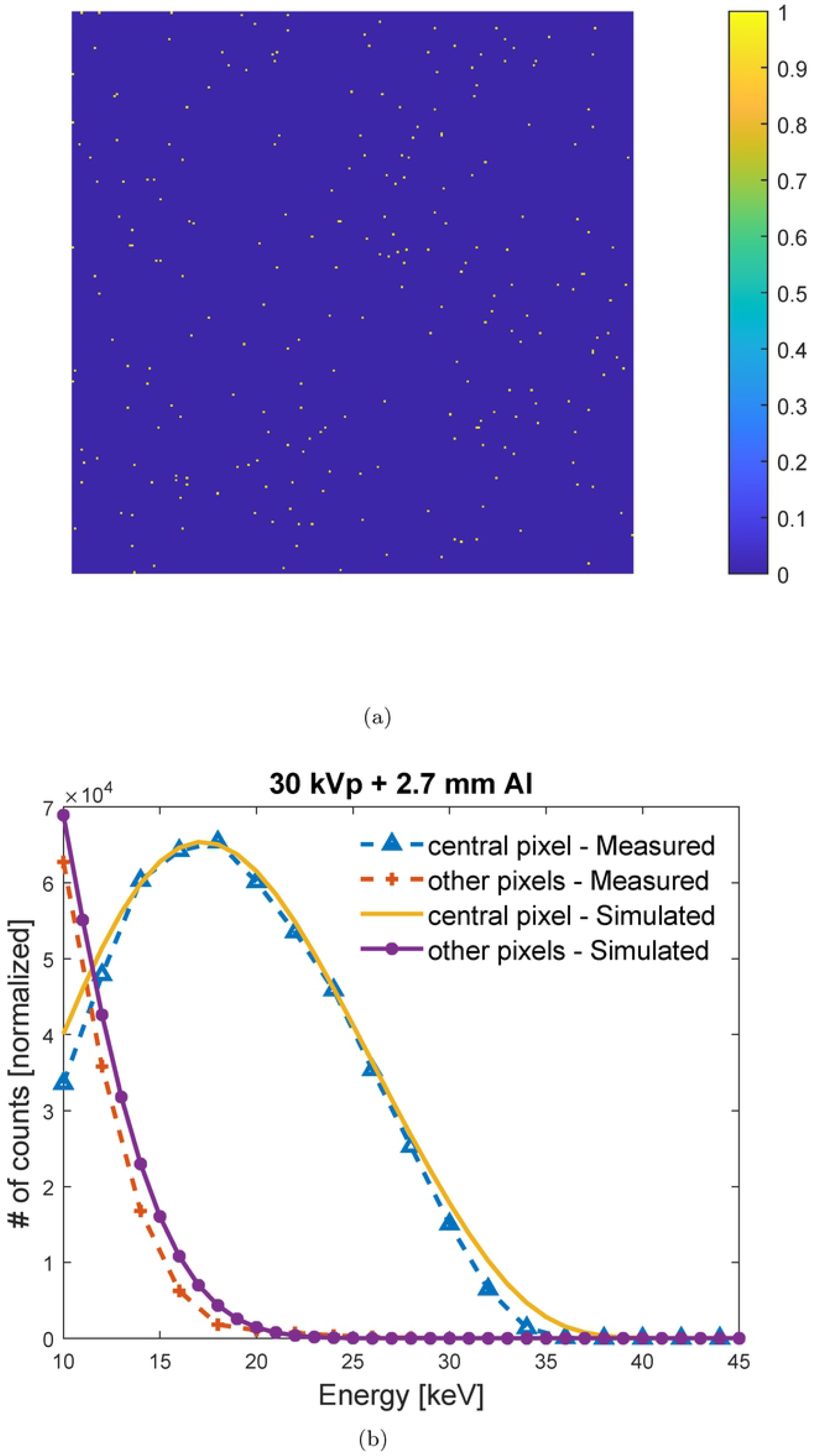
(a) Sample image of single events interactions recorded at energy threshold of 15 keV. (b) Comparison between the experimental and simulated spectra estimated using the single events analysis method when using 11 *µ*m and 2.1 keV for *r*_0_ and *σ*_*e*_, respectively.

## Conclusion

In this study, we presented an extension to the PcTK, dedicated to the simulation of a realistic GaAs PCD. The extended version was evaluated by comparison between simulated and measured spectra. The first validation was by comparison between simulated and measured EDXRD spectra from a 109Cd radionuclide source and was also aimed at obtaining the optimum model parameters for simulating the LAMBDA 60 K module planar detector used in this study. The second validation involved evaluating the performance of the GaAs version of the PcTK with polychromatic radiation used in conventional x-ray imaging systems. The third validation experiment was to validate the spatio-energetic model of the GaAs version of the PcTK using single events analysis. Overall, the software provided a good agreement between simulated and experimental data, validating the accuracy of the GaAs model. The PcTK can now be used to design, study and optimize the parameters affecting the image quality of breast CT, DBT and mammography imaging systems with GaAs-based photon counting detectors.

